# Toward Site-Specific Characterization of Structural Perturbations on Glycosylated Fc Using NMR at Natural Abundance

**DOI:** 10.64898/2025.12.03.692071

**Authors:** Béatrice Vibert, Sarah Nguyen, Faustine Henot, Camille Doyen, Oscar Hernandez-Alba, Sarah Cianférani, Séverine Clavier, Oriane Frances, Jérôme Boisbouvier

## Abstract

Monoclonal antibodies (mAbs) are leading therapeutic agents due to their high specificity and limited side effects. Ensuring their structural integrity under stress and maintaining batch consistency require robust quality control. Methyl 2D NMR has emerged as a powerful tool to probe mAb structure at natural isotopic abundance, enabling spectral fingerprint comparisons across production batches to detect subtle structural changes. However, extracting atomic-level structural information requires assignment of methyl resonances to their amino acids. While such assignments are available for several antigen-binding fragments (Fabs), no comprehensive assignment has been reported for the crystallisable fragment (Fc). In this study, we present the methyl group assignment of the 50-kDa Fc fragment of an immunoglobulin G1 (IgG1) antibody. Using cell-free expression, strategic isotopic labelling, and high-quality 2D and 3D NMR experiments, we successfully assigned 94% of methyl resonances of a non-glycosylated Fc. Given that therapeutic mAbs are typically produced in Chinese Hamster Ovary (CHO) cells, we transferred this assignment to the methyl spectrum of a glycosylated Fc fragment obtained by the enzymatic cleavage of a CHO-produced mAb at natural abundance, achieving 83% assignment coverage. This assignment was then used to investigate the impact of methionine oxidation on Fc structure at atomic resolution using NMR. The methyl group assignment transforms 2D methyl NMR fingerprinting into a powerful tool for quality control. It enables the direct comparison of spectra acquired on mAbs produced at natural abundance, allowing the detection and localisation of chemical modifications and structural changes without the need for isotopic labelling. This approach offers a robust solution for monitoring the structural integrity of therapeutic antibodies throughout development and manufacturing.

## Introduction

Monoclonal antibodies (mAbs) currently represent the most prominent class of pharmaceutical biomolecules, with a record of 13 mAbs approved by the Food and Drug Administration (FDA) in 2024 (de la Torre and Albericio 2025; Martins et al. 2025). These biomolecules are engineered to target antigens with high specificity and minimal side effects (Kothari et al. 2024). Rigorous quality control is therefore essential, not only during development phases to monitor degradation under stress conditions, but also throughout manufacturing to ensure batch-to-batch consistency. However, their large size, structural complexity and elaborate production in mammalian cells make them significantly more challenging to characterize than small-molecule therapeutics.

mAbs are 150-kDa proteins, consisting of three 50-kDa fragments: the crystallisable fragment (Fc), which mediates interactions with immune system receptors, and the two antigen-binding fragments (Fabs). Both are 50-kDa multi-chain proteins rich in disulfide bridges, with the Fab being a heterodimer and the Fc a homodimer. Each antibody comprises two identical heavy chains (H) and two identical light chains (L), organized into variable (V) and constant (C) domains: V_L_, C_L_, V_H_, and C_H_1 for the Fab; C_H_2 and C_H_3 for the Fc.

Methyl groups are well-suited NMR probes for investigating the structure of high-molecular-weight proteins (Tugarinov and Kay 2005; Kerfah et al. 2015) and, in recent years, methyl-based NMR has emerged as a complementary tool to established biophysical techniques commonly used for mAb quality control, providing atomic-level structural fingerprints of mAbs (Brinson et al. 2018; Marino et al. 2015; Cerofolini et al. 2023; Arbogast et al. 2017). The favourable properties of methyl groups arise from the presence of three equivalent protons, their fast rotation, and their location at the end of side chains. Moreover, methylated amino acids account for approximatively 30 to 40% of protein residues and are evenly distributed throughout the protein structure, providing a robust network of structural probes. Two-dimensional ^1^H-^13^C methyl spectra serve as structural fingerprints and can be acquired across a wide range of proteins, with or without isotopic labelling (Rößler et al. 2020). These favourable properties make methyl groups particularly well-suited for probing the structure and dynamics of complex biomolecules, such as monoclonal antibodies. While these 2D spectral fingerprints are already used to compare mAb batches under stress conditions at natural isotopic abundance (Solomon et al. 2023), obtaining residue-specific information requires assigning each methyl signal to its corresponding amino acid. This would enable the precise localisation of stress-affected residues within the 3D structure of the antibody or its individual fragments. To study these fragments by NMR and assign their methyl signals, isotopic labelling and high-quality 3D NMR experiments are required. Such studies have focused on immunoglobulin G1 (IgG1) Fabs, including model NISTmAb (Ghasriani et al. 2022), adalimumab (Sarker and Aubin 2024), trastuzumab (Gagné et al. 2024), anti-LAMP1 mAb (Henot et al. 2025), and ipilimumab (Henot et al. 2025). Labelled samples were produced in *E. coli* (Ghasriani et al. 2022), cell-free system (Giraud et al. 2024), or yeast (Gagné et al. 2023), with numerous innovative and well-designed labelling strategies. Various constructs were also developed to facilitate Fab production and assignment. Typically, backbone amide resonances were assigned first (Giraud et al. 2024), followed by the transfer of assignments to intra-residue side chain methyl groups using 3D HCC TOCSY or COSY experiments. NOESY experiments were performed in some cases to complete the assignment of methyl groups. Unlike the Fab fragment, which varies between antibodies, the Fc fragment is conserved across IgG1s. However, it contains glycans attached to asparagine N297 in the C_H_2 domain of both chains. These glycans, as post-translational modifications, are absent in Fc fragments expressed in bacterial derived systems. NMR studies have been conducted on both non-glycosylated Fc produced in *E. coli* and glycosylated Fc obtained from mAbs produced in CHO cells, resulting in NH backbone assignments (Liu et al. 2007a; Yagi et al. 2015). Nonetheless, even if it is a prerequisite for site-specific NMR study at natural abundance, the methyl group assignment was not achieved in these studies. Hence, only the methyl group assignment of the reduced C_H_3 domain is currently available (Liu et al. 2007b).

In this work, we present a near-complete assignment of methyl group signals of the Fc fragment of an IgG1 antibody. In pharmaceutical industry, mAbs are produced in CHO cells thus undergo glycosylation, a post-translational modification essential for their stability and biological activity. However, isotopically labelled protein production is more feasible and adaptable in bacterial derived expression systems than in mammalian cells. Therefore, we first assigned the methyl groups of the non-glycosylated IgG1 Fc fragment. Samples were produced using a cell-free system based on *E. coli* extracts, allowing the production of correctly folded Fc fragment without the need for refolding steps, enabled by the addition of a disulfide bond isomerase in the cell-free reaction mixture. Using available backbone NMR data (Liu et al. 2007a) and a series of 2D and 3D NMR experiments acquired on various isotopically labelled samples, a near-complete assignment of side-chain methyl groups was achieved. This assignment was then transferred to the 2D methyl spectrum of a glycosylated Fc fragment from a CHO cell-produced mAb at natural abundance, and the signals showing significant chemical shift differences between the two Fc constructs were highlighted. We leveraged this methyl group assignment, covering 83% of the methylated residues of the glycosylated Fc, to study stress-induced modifications in an anti-lysosomal-associated membrane protein 1 (anti-LAMP1) mAb, produced by Sanofi for research purposes. Using Fc fragments obtained directly from therapeutic mAb batches, we investigated the impact of methionine oxidation on Fc structure at atomic resolution using NMR.

## Materials and methods

### Preparation of the non-glycosylated Fc fragment for NMR assignment

Plasmid preparation: the DNA sequence of an IgG1 Fc (residues D221 to G446) was cloned in pIVEX 2.3d plasmid between NcoI and SmaI restriction enzymes. The C-terminal region of the plasmid contained a TEV protease cleavage site (ENLYFQG) followed by a 6-His tag (see supplementary information for the detailed sequence). The plasmid was transformed into TOP10 *E. coli* strain grown in 500 mL of LB medium with 100 µg/mL of ampicillin. The plasmid was then purified using NucleoBond Xtra Maxi Plus kit (Macherey-Nagel) and the final plasmid concentration was adjusted around 1 mg/mL of DNA in RNase-free water.

DsbC production: a pET-28a plasmid containing DsbC DNA sequence with a N-terminal 6-His tag followed by a TEV protease cleavage site was transformed into *E. coli* BL21 (DE3) strain. Bacteria were grown in TB medium and overexpression of DsbC was induced using 1 mM of IPTG. After 16 h at 20 °C, cells were collected and lysed using sonication. After centrifugation, the supernatant was recovered and DsbC was purified using a Ni-NTA affinity column followed by size-exclusion chromatography (SEC) separation and finally concentrated to about 200 µM in 50 mM HEPES (Euromedex), 150 mM NaCl and 5% glycerol at pH 7.5.

Cell-free expression: cell-free S30 lysates were prepared using an *E. coli* BL21 (DE3) strain following the previously published protocol (Imbert et al. 2021). DTT was omitted in the dialysis buffer to prevent a possible reduction of disulfide bridges of the Fc fragment. All cell-free reactions were performed at 30 °C with a gentle shaking (20 rpm) using a hybridization oven (Techne). To optimize betaine, magnesium and DsbC concentrations, 50 µL cell-free syntheses of the Fc fragment were undertaken in 1.5 mL Eppendorf tubes. Reactions were stopped after 3 h and analysed with Western blots (Bio-Rad kits) using anti-his-tag antibody. Fc production was performed in continuous exchange mode cell-free (CECF) using GeBAflex dialysis devices (Euromedex) filled with 3 mL of reaction mixture and placed in 25 mL centrifugation tubes containing 15 mL of dialysis mixture. Standard cell-free components including nucleotides, HEPES, folinic acid, cyclic AMP, spermidine, NH_4_OAc and creatine phosphate were added in both the reaction and the dialysis mixtures as described in a previously published protocol (Imbert et al. 2021). Betaine, magnesium, and DsbC were added at their respective optimal concentrations: 15 mM, 14 mM and 14 µM. The 20 amino acids were added at a final concentration of 1 mM each. To control the redox potential of the reaction, both reduced and oxidized glutathione was used at concentrations of 0.9 and 3.6 mM, respectively, according to previously published protocols (Yin and Swartz 2004; Giraud et al. 2024). The reaction mixture was completed with creatine kinase, T7 RNA polymerase, tRNA, DsbC, S30 bacterial extract, and 16 µg/mL of DNA plasmid and the reaction was run for 16 h.

Isotopic labelling: to produce uniformly ^13^C, ^15^N labelled samples, the amino acid mixture was replaced with an algal-derived amino acid mix (Isogro® ^13^C, ^15^N powder growth medium) hydrolysed as described previously (Imbert et al. 2021), supplemented with unlabelled tryptophan and cysteine at a final concentration of 1 mM each. Fc samples specifically labelled on methyl groups were obtained by adding each of the 20 amino acids with the desired labelling directly to the cell-free reaction mixture. For Ile-δ_1_, Leu-δ_1_/δ_2_, and Val-γ_1_/γ_2_, labelled precursors were added in place of the corresponding amino acids (Gans et al. 2013). 2-keto-3-(CD_3_-methyl)-3,4,4-(D_3_)-5-(^13^C)-pentanoate was used to selectively label isoleucine at the δ_1_ position (Henot et al. 2025). α-ketoisovaleric acid 3-D_1_, containing two ^13^CH_3_ methyl groups, was used to label valine at both γ_1_ and γ_2_ positions simultaneously (Goto et al. 1999). A variant with one ^13^CH_3_ and one ^12^CD_3_ methyl group was employed to achieve 50% labelling at γ_1_ and 50% at γ_2_, without stereospecificity (Tugarinov and Kay 2004). Uniformly ^13^C-labelled α-ketoisocaproic acid was used to label leucine at both δ_1_ and δ_2_ positions. The labelling scheme used for each sample is presented in Table S1.

Purification: CECF reaction mixture was centrifuged for 20 min at 14,000 rpm. Supernatant was diluted by a factor of 4 in PBS 1X (Euromedex) at pH 7.4 and purified using a 1 mL CaptureSelect FcXP (Thermo Scientific) affinity chromatography column. For 3 mL of cell-free reaction mixture, the column was loaded with 12 mL of diluted supernatant and washed with 15 mL of PBS 1X at pH 7.4. The Fc was eluted with a 5 mL gradient from 0 to 100% of 20 mM acetic acid at pH 5.0 in PBS, followed by 10 mL of 20 mM acetic acid at pH 4.0. 20 µL of 1 M Tris base (Euromedex) buffer at pH 8.0 were immediately added to each 1 mL elution fraction. The Fc fragment was buffer-exchanged and concentrated in the NMR buffer containing 10 mM NaOAc at pH 5.0, EDTA-free cOmplete protease inhibitor cocktail (Roche) and 10 or 100% of D_2_O using a centrifugal filtration unit (Amicon® 10 kDa MWCO). Final concentrations ranging from 0.16 mM to 1.6 mM were achieved, depending on the labelling scheme (see Table S1 for the exact concentration of each sample). The final samples were placed in either 4 mm Shigemi tubes or 3 mm standard NMR tubes. See Table S1 for samples’ details and Fig. S1 for Fc quality control.

### Preparation of glycosylated Fc fragments from anti-LAMP1 mAb for oxidation study

Oxidation of anti-LAMP1 mAb: oxidation was carried out by incubating a 30 mg/mL solution of anti-LAMP1 mAb overnight at 37 °C with 2.4% (v/v) of tert-Butyl hydroperoxide (tBHP). The reaction was stopped by performing a buffer exchange into Dulbecco’s phosphate-buffered saline (DPBS) using Zeba™ Spin desalting columns. This exchange procedure was also applied to the non-oxidized sample. Preparation of Fc samples for NMR acquisition: enzymatic cleavage of both oxidized and non-oxidized mAbs was performed using the FabALACTICA^TM^ Fab kit Maxispin (Genovis). 600 µL of mAb were added to 4 mL of reaction buffer (150 mM sodium phosphate, pH 6.7), followed by an overnight incubation at room temperature with end-over-end mixing. After elution, the processed material was loaded onto a CaptureSelect^TM^ Kappa XP column, and the Fc fragment was collected in the flow-through. A second purification step was performed using Protein A affinity chromatography. The Fc fragment was eluted with a glycine solution at pH 2.7 and subsequently neutralized by addition of a Tris solution at pH 9. The Fc fragment was then buffer-exchanged in 10 mM NaOAc at pH 5.0 in 100% of D_2_O, and concentrated to approximatively 10 mg/mL using a centrifugal filtration unit with a 10-kDa cut-off. Samples were transferred into 3 mm NMR tubes.

### Mass spectrometry analysis

Non-denaturing middle-level mass spectrometry analysis: mass spectrometry analysis of subunits was performed using a ZenoTOF 7600 (SCIEX) mass spectrometer, hyphenated with liquid chromatography (LC). After FabRICATOR (IdeS) digestion of anti-LAMP1 mAb, F(ab’)_2_ and Fc subunits were separated by SEC using an isocratic flow rate of 250 µL/min of 100 mM ammonium acetate. Non-denaturing MS conditions included a scan range of 1000-10000 m/z, a spray voltage of 4500 V, a source temperature of 350 °C, a declustering potential of 100 V, a collision energy of 4 V, and a gas pressure of 35 psi to ensure efficient ion transmission. MS data were processed using SCIEX OS 3.0 and Protein Metrics softwares.

Peptide mapping: prior to analysis, samples were enzymatically digested with trypsin, following denaturation, reduction, and alkylation steps. LC-MS/MS experiments were conducted on a Q-Exactive Plus (Thermo Scientific) mass spectrometer. Peptides were separated using a typical reversed-phase liquid chromatography (RPLC) bottom-up gradient prior to analysis on a C18 column over a 110-min run. Data were processed using Protein Metrics.

### NMR spectroscopy

Backbone amide assignment of the non-glycosylated Fc fragment: to identify each NH backbone signal (Liu et al. 2007a), NMR data were acquired on a 0.3 mM U-[^15^N, ^13^C] non-glycosylated Fc sample using Bruker spectrometers equipped with cryogenic probes operating at ^1^H frequencies of 600 to 950 MHz. ^1^H-^15^N BEST-TROSY 2D experiments (Lescop et al. 2007; Pervushin et al. 1997) were recorded at temperatures ranging from 303 K to 323 K to enable assignment transfer from published data to acquired spectra. The assignment was completed using 3D BEST-TROSY HNCO and HNCA triple resonance experiments (Lescop et al. 2007) acquired at 323 K.

Methyl group assignment of the non-glycosylated Fc fragment: all NMR experiments used to assign methyl group resonances of the non-glycosylated Fc fragment were recorded at 323 K on Bruker spectrometers equipped with cryogenic probes operating at ^1^H frequencies of 600 to 950 MHz. 2D ^1^H-^13^C methyl SOFAST-HMQC experiments were recorded to identify each methyl type using samples labelled as follows: Ala-[^13^C^1^H_3_]^β^, Ile-[U-^2^H; [^13^C^1^H_3_]^δ1^] and Thr-[U-^2^H; [^13^C^1^H_3_]^γ^], Met-[^13^C^1^H_3_]^ε^ and Val-[U-^2^H; [^13^C^1^H_3_]^γ1/γ2^], and Leu-[U-^13^C; [^13^C^1^H_3_]^δ1,δ2^] (Schanda and Brutscher 2005). Methyl group assignment was initiated using a 3D hCCH TOCSY experiment acquired on a 0.7 mM U-[^15^N, ^13^C] Fc sample with a transfer delay of 21 ms, an interscan delay of 1.0 s, and acquisition time of 8.3 ms in the ^13^C indirect dimensions and 70 ms in ^1^H direct dimension (Kay et al. 1993). To complete the assignment of methyl group resonances, 3D CCH HMQC-NOESY-HMQC NMR experiments (Tugarinov et. al 2005) were acquired on three different Fc samples produced in H_2_O with protonated amino acids and buffer-exchanged in D_2_O for NMR acquisition, with the following labelling schemes: *i)* Ala-[^13^C^1^H_3_]^β^, Ile-[U-^2^H; [^13^C^1^H_3_]^δ1^], Met-[^13^C^1^H_3_]^ξ^, Thr-[U-^2^H; [^13^C^1^H_3_]^γ^], and Val-[U-^2^H; [^13^C^1^H_3_]^γ1/γ2^]; *ii)* Leu-[U-^13^C; [^13^C^1^H_3_]^δ1,δ2^]; and *iii)* Val-[U-^2^H; [^13^C^1^H_3_]^γ1,γ2^]. For the first sample, the NOE mixing time was 125 ms, the interscan delay was 0.96 s, and the acquisition time was 20 ms in the ^13^C indirect dimensions and 60 ms in the ^1^H direct dimension. For the two other samples, the NOE mixing time was 150 ms, the interscan delay was 0.79 s and the acquisition time was 14 ms in the ^13^C indirect dimensions and 60 ms in the ^1^H direct dimension. See Table S1 for details on NMR experiments and samples used for methyl group assignment. All NMR experiments detailed above were acquired using NMRlib (Favier and Brutscher 2019).

For glycosylated Fc fragment at natural abundance before and after oxidation, 2D ^1^H-^13^C ALSOFAST-HMQC (Arbogast et al. 2018; Mueller 2008) experiments were recorded on a spectrometer operating at a ^1^H frequency of 800 MHz, at 313 K. Acquisition times of 62 ms in the ^1^H dimension and 8 or 21 ms in the ^13^C dimension were used, and the interscan delay was set at 0.6 s.

All data were processed using NMRPipe (Delaglio et al. 1995) and Topspin 4.1.4 (Bruker), and analysed using CcpNmr 3.2.2 (Vranker et al. 2005).

## Results and discussion

### Methyl group resonances assignment of the Fc fragment produced in a cell-free system

To enable high-resolution NMR study of the Fc fragment, samples uniformly and specifically enriched with ^13^C and ^15^N were produced *in vitro* using a cell-free system. As previously described (Giraud et al. 2024), the cell-free reaction mixture can be supplemented with the disulfide bond isomerase DsbC, and its redox potential can be controlled to ensure the formation of the native disulfide bridges topology of mAb fragments. After optimization of expression and purification conditions, a yield of 0.4 mg of Fc per mL of cell-free reaction was achieved at natural isotopic abundance. The open nature of the cell-free system also enabled tailored isotopic labelling strategies, including residue-specific and stereospecific labelling.

To assign the methyl group resonances of the 50-kDa Fc fragment, we employed a multi-step strategy combining backbone-to-methyl transfer techniques via 3D TOCSY and COSY experiments with NOESY-based spatial correlation (Liu et al. 2007; Sarker and Aubin 2024; Gagné et al. 2024; Ghasriani et al. 2022; Henot et al. 2025). The assignment of backbone amide resonances of the non-glycosylated human IgG1 Fc had already been undertaken by Liu et al., resulting in the assignment of 93% of C^α^ and 92% of C^β^ resonances from methylated amino acids (Liu et al. 2007a). To validate the backbone assignment on the 2D ^1^H-^15^N HSQC spectrum acquired on the Fc fragment produced in a cell-free system, 3D BEST-TROSY HNCO and HNCA experiments were acquired on a 0.3 mM U-[^15^N, ^13^C] Fc sample. As a result, NH backbone resonances were assigned to their corresponding amino acids for 90% of the Fc’s methylated amino acids (Fig. S2).

To facilitate the methyl group assignment, 2D ^1^H-^13^C spectra were acquired on four Fc samples specifically labelled on Ala-β, Ile-δ_1_ and Thr-γ, Met-ε and Val-γ_1,2_, and Leu-δ_1,2_. An hCCH TOCSY experiment was acquired on a 0.7 mM U-[^15^N, ^13^C] Fc sample, enabling the connection of signals from the 2D ^1^H-^13^C SOFAST-HMQC methyl spectrum to the side-chain carbon resonances of the corresponding methylated residues (Fig. 1a and Fig. 1b). By comparing the retrieved C^α^ and C^β^ resonances with published data (Liu et al. 2007a), 42% of the methylated residues were assigned, including 71% of alanines, 75% of isoleucines, 42% of leucines, 44% of threonines, and 26% of valines with at least one assigned methyl group. Despite the high coverage of backbone assignments, the transfer to methyl signals was limited by lack of deuteration, signal overlap, and discrepancies with published chemical shifts, which made assignment transfer ambiguous in crowded regions of the methyl spectrum.

**Fig. 1.**
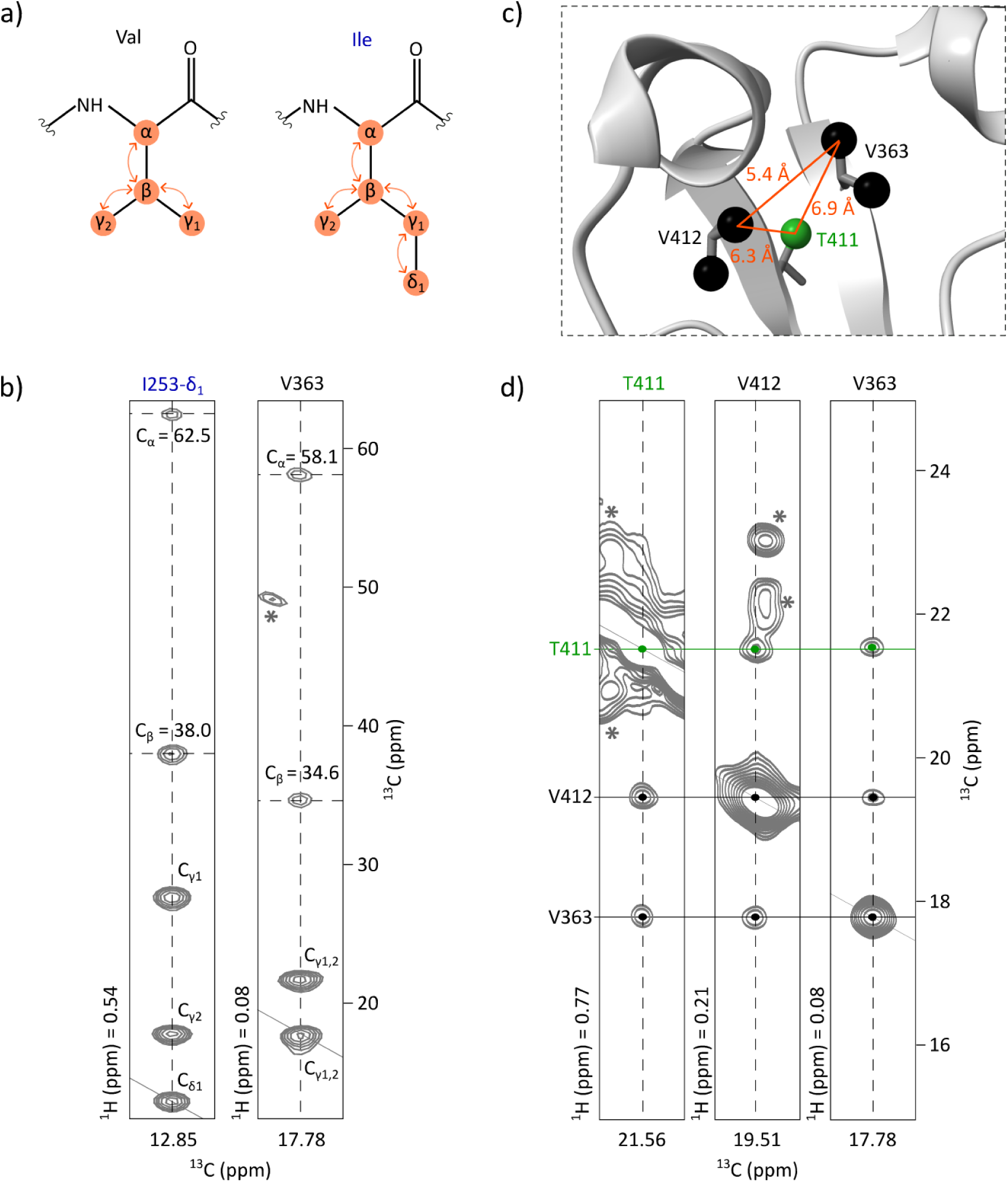
Methyl group assignment strategy for the non-glycosylated Fc fragment. **a)** Scheme of the magnetization transfer (represented with orange arrows) occurring during TOCSY mixing time in valine and isoleucine residues. **b)** Assignment transfer from the backbone to methyl groups using an hCCH-TOCSY experiment acquired on a U-[^15^N, ^13^C] sample. 2D extracts are displayed correlating previously assigned C^α^ and C^β^ resonances to the methyl resonances of V363 and I253. Asterisks indicate signals outside of the considered plans. **c)** Zoom on V412, V363, and T411 extracted from the 3D structure of the non-glycosylated Fc fragment (PDB: 3DNK) showing Ala-β, Ile-δ_1_, Val-γ_1_, Val-γ_2_, Thr-γ and Met-ε methyl groups. **d)** 2D extracts from a 3D HMQC-NOESY-HMQC experiment acquired on an Fc fragment labelled on Ala-[^13^C^1^H_3_]^β^, Ile-[U-^2^H; [^13^C^1^H_3_]^δ1^], Met-[^13^C^1^H_3_]^ξ^, Thr-[U-^2^H; [^13^C^1^H_3_]^γ^], and Val-[U-^2^H; [^13^C^1^H_3_]^γ1/γ2^] showing NOE cross-peaks between V363, V412, and T411. Asterisks indicate signals outside of the considered plans

These assigned methyl resonances were used as a starting point for the analysis of a 3D methyl-methyl CCH NOESY experiment acquired on an Fc sample labelled as follows: Ala-[^13^C^1^H_3_]^β^, Ile-[U-^2^H; [^13^C^1^H_3_]^δ1^], Met-[^13^C^1^H_3_]^ξ^, Thr-[U-^2^H; [^13^C^1^H_3_]^γ^] and Val-[U-^2^H; [^13^C^1^H_3_]^γ1/γ2^]. NOE cross-peaks revealed spatial proximity (≤ 10 Å) between methyl groups of alanines, isoleucines-δ_1_, methionines, threonines, and valines that are close in space. Signals were manually analysed using the 3D structure of a non-glycosylated Fc fragment (PDB: 3DNK), along with the amino acid types of each methyl signal previously determined from specifically labelled Fc samples (Fig. 1c and Fig. 1d). By combining these data with the previously assigned methyl resonances, assignment was achieved up to 100% for alanines, isoleucines, and methionines, and at 69% for threonines. However, no leucines and only 13% of additional valines were assigned. This was mainly due to the labelling scheme: leucines were not labelled and half of the valines were labelled on γ_1_ and the other half on γ_2_, decreasing the expected NOE cross-peak intensity by a factor of four for valine-valine pairs and by a factor of two for valine-non-valine pairs. To further complete the assignment, two additional samples were prepared, labelled respectively on Leu-[U-^13^C; [^13^C^1^H_3_]^δ1,δ2^] and Val-[U-^2^H; [^13^C^1^H_3_]^γ1,γ2^], and similar NOESY experiments were acquired. These additional datasets not only enabled the correlation of the two methyl signals for each leucine and valine residue, but also confirmed previously ambiguous amino acid assignments (Fig. S3). Furthermore, they enabled the assignment of additional methyl group resonances for valines and leucines, as some of these residues are clustered within the Fc structure. Ultimately, at least one methyl resonance was assigned for 100% of valine and 95% of leucine residues. The complete assignment strategy is summarized in Fig. S4.

By combining all these experimental data and labelling strategies, we achieved a remarkably high assignment coverage of the methyl resonances of the non-glycosylated Fc fragment produced using a cell-free system. Specifically, 94% of the methyl signals were successfully assigned, representing a comprehensive mapping of the structural probes within this 50-kDa fragment (Fig. 2). Only three threonine and one leucine residues remained unassigned. This near-complete assignment provides a robust foundation for structural investigations of the Fc region. It not only enables high-resolution analysis of the non-glycosylated Fc form but also represents a strategic step toward extending investigations to more physiologically relevant systems. To achieve this, it is essential to transfer the methyl resonance assignment to glycosylated Fc fragments produced in CHO cells at natural isotopic abundance. This would enable direct monitoring of structural integrity and the detection of stress-induced modifications directly on therapeutic mAb batches during development and manufacturing.

**Fig. 2.**
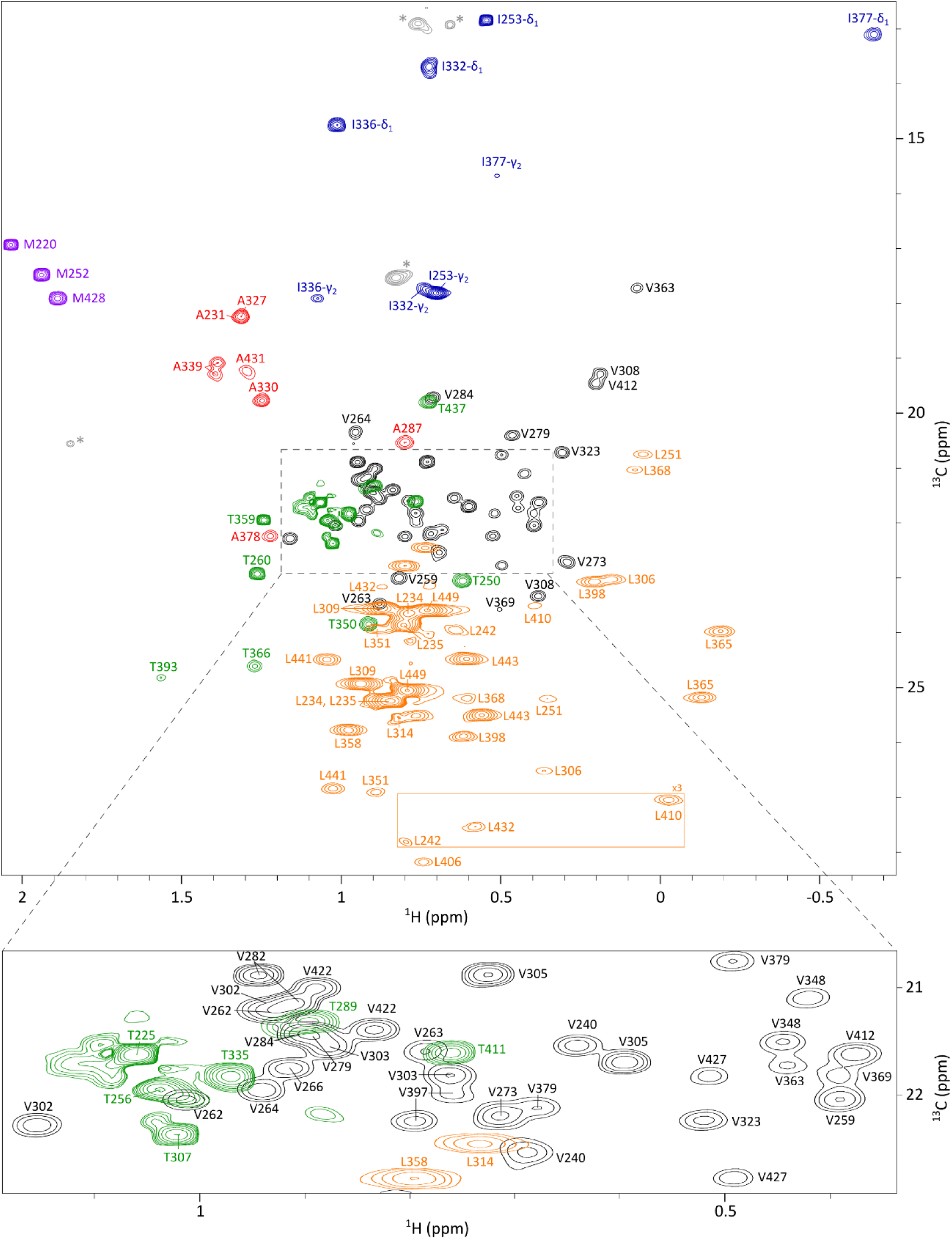
Assigned 2D ^1^H-^13^C SOFAST HMQC spectrum of the non-glycosylated Fc fragment produced using a cell-free system. This spectrum is a superimposition of four spectra of the Fc fragment labelled on Ala-[^13^C^1^H_3_]^β^, Ile-[^13^C^1^H_3_]^δ1^ and Thr-[^13^C^1^H_3_]^γ^, Leu-[^13^C^1^H_3_]^δ1,δ2^, and Met-[^13^C^1^H_3_]^ε^ and Val-[^13^C^1^H_3_]^γ1/γ2^ together with a zoom on Ile-γ_2_ region from a spectrum acquired on a U-[^15^N, ^13^C] Fc sample. Alanine, isoleucine, leucine, methionine, threonine, and valine resonances are depicted respectively in red, blue, orange, purple, green, and black. Signals depicted in grey and annotated with an asterisk correspond to impurities. Each assigned signal is annotated with the corresponding residue number. The contour level of signals in the orange rectangle has been multiplied by a factor of 3

### Assignment transfer to a glycosylated Fc fragment produced in CHO cells

Building on the methyl assignment of the non-glycosylated Fc fragment, we extended our analysis to glycosylated Fc fragments at natural abundance derived from monoclonal antibodies produced in CHO cells, conditions that closely reflect industrial production processes. Since all IgG1 antibodies share the same Fc fragment, the assignment obtained for the Fc produced in a cell-free system can, in principle, be transferred to any IgG1. However, glycosylation being attached to the C_H_2 domain, chemical shift perturbations may affect methyl signals for residues in this region, highlighting the need for a careful assignment transfer. To ensure high-quality methyl spectra suitable for assignment transfer and atomic-scale analysis, the 50-kDa Fc fragment was used instead of the full 150-kDa monoclonal antibody. This choice enabled an increase of the acquisition time in the carbon dimension from 8 to 21 ms, improving spectral resolution while maintaining a good signal-to-noise ratio (Fig. 3). Interestingly, all Fc methyl signals overlapped with those of the mAb, further supporting the use on this divide-and-conquer strategy.

**Fig. 3.**
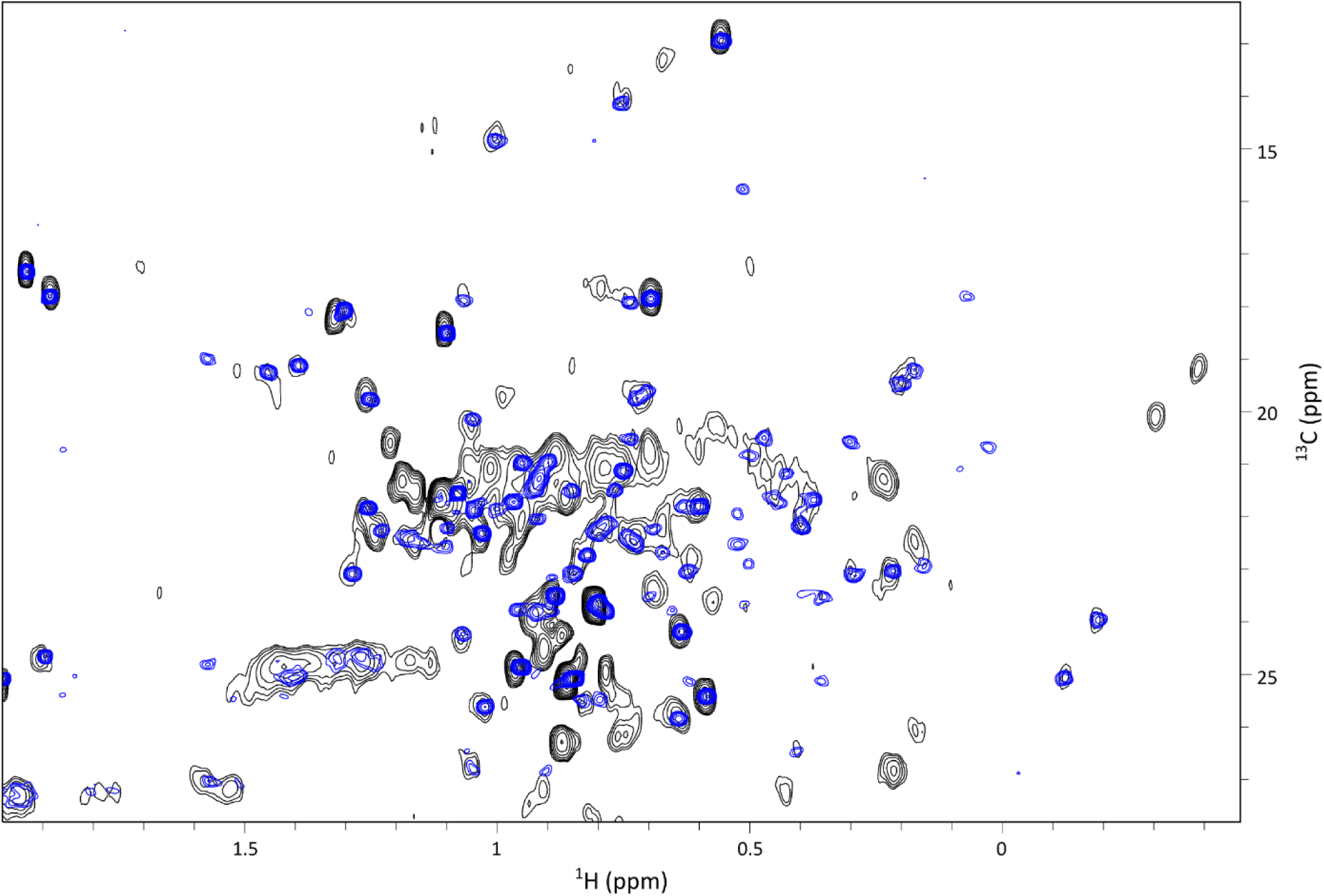
Superimposition of ^1^H-^13^C SOFAST methyl spectra of anti-LAMP1 mAb in black with its Fc fragment in blue. To generate the Fc fragment, the mAb was cleaved above the hinge between T223 and H224 using an IgdE enzyme (FabALACTICA^TM^, Genovis). Spectra were acquired on samples at natural abundance on an 800 MHz spectrometer, with an acquisition time in carbon dimension of 8 ms for the mAb and 21 ms for the Fc

The methyl group assignment of the crystallisable fragment was transferred from the isotopically labelled, non-glycosylated Fc produced in a cell-free system to the glycosylated Fc obtained by enzymatic cleavage of a mAb produced in CHO cells (Fig. S5). Overlaying the 2D methyl spectra of both Fc constructs enabled successful assignment transfer for 86% of alanines, 100% of isoleucines, 94% of leucines, 100% of methionines, 77% of threonines, and 83% of valines, resulting in an overall assignment of 83% of methylated residues in the glycosylated Fc spectrum (Fig. 4). The main limitation arises from the difficulty of transferring assignments in crowded regions of the spectrum, further complicated by the low intensity of some methyl signals in the spectrum acquired on the Fc sample at natural abundance. This low intensity can be partly due to the glycan heterogeneity, decreasing the intensity of the signals impacted by the glycans. Nevertheless, all well-resolved signals in non-crowded regions were successfully assigned.

**Fig. 4.**
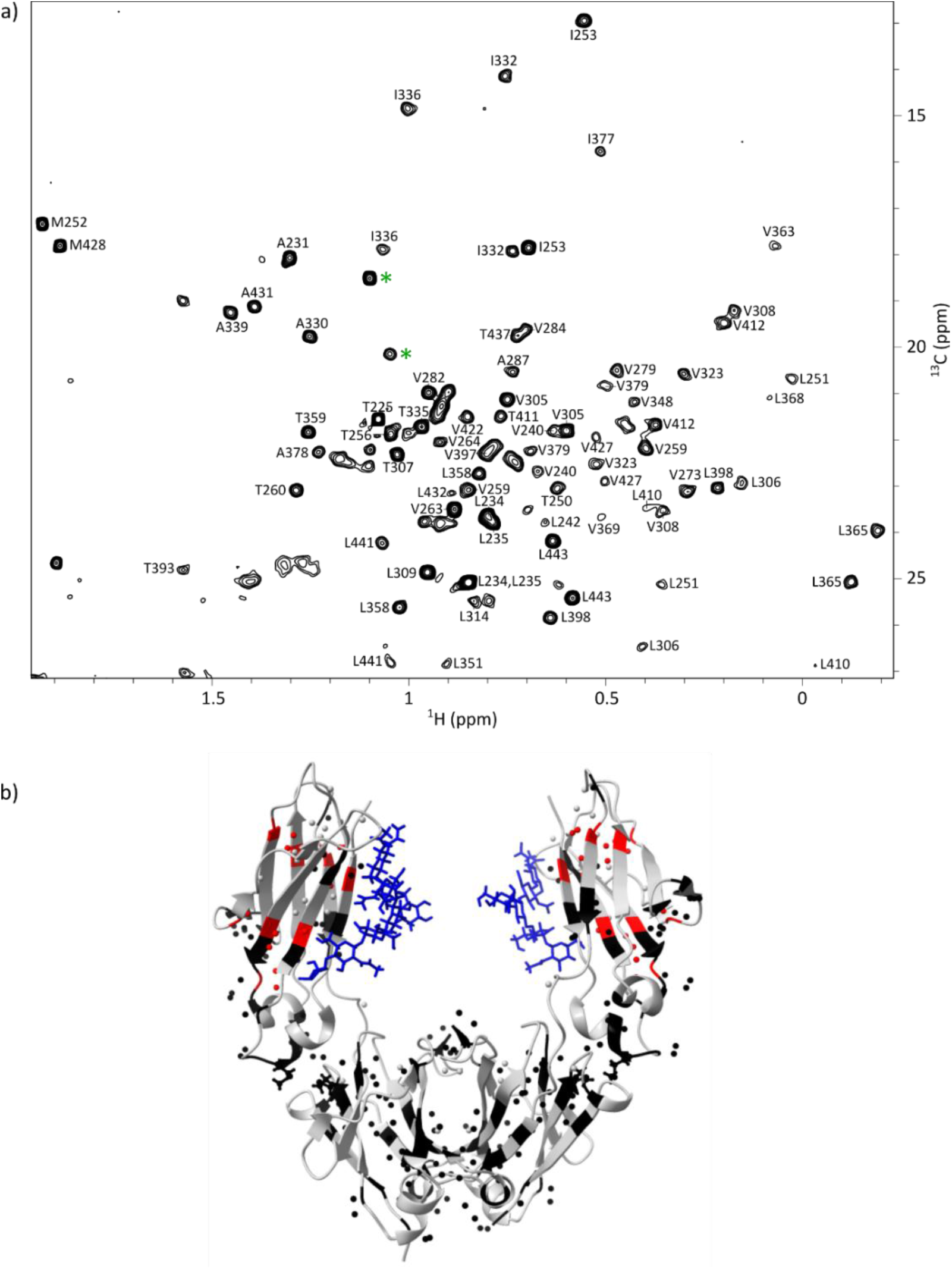
**a)** Assigned ^1^H-^13^C ALSOFAST methyl spectrum of the glycosylated Fc fragment from anti-LAMP1 mAb produced in CHO cells at natural abundance. The two signals annotated with a green asterisk correspond to methyl groups of *N*-acetylglucosamine (GlcNAc) moieties from the glycans. **b)** Fc structure (PBD: 3JII) showing the residues with important shift of methyl resonances between the ^1^H-^13^C spectrum of the non-glycosylated Fc fragment produced in a cell-free system and the spectrum of the glycosylated Fc fragment from anti-LAMP1 produced in CHO cells. Glycans are represented in blue. Methyl groups are represented by spheres. Non-methylated amino acids and non-assigned methylated amino acids are in grey, assigned methylated amino acids are in black when CSP < 0.03 ppm and in red when CSP ≥ 0.03 ppm. CSP values between glycosylated and non-glycosylated Fc were calculated using the following formula: 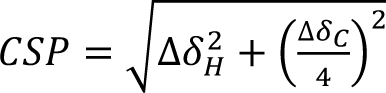. For residues with two methyl groups (isoleucines, leucines, and valines), the average CSP was used when both signals were assigned

Interestingly, two additional signals were detected in the glycosylated Fc spectrum that were absent in the non-glycosylated one (signals marked with asterisks in Fig. 4). To investigate their origin, we acquired a methyl spectrum of a deglycosylated Fc fragment, obtained from a CHO-produced mAb, and compared it to the spectrum of its glycosylated counterpart. While most of the peaks were either perfectly superimposed or slightly shifted due to structural changes upon deglycosylation, the two signals of interest completely disappeared, confirming their assignment to the methyl groups of *N*-acetylglucosamine moieties from the glycans (Fig. S6).

To further characterise the structural differences between the two Fc constructs, chemical shift perturbations (CSPs) were calculated for each assigned methyl signal (Fig. S7). As expected and illustrated in Fig. 4, the largest perturbations were observed for residues located in the C_H_2 domain, where glycans are attached. Despite these local changes, the successful transfer of methyl assignments by the simple overlay of the two 2D spectra demonstrates the feasibility of atomic-resolution NMR analysis of glycosylated IgG1 antibodies at natural abundance. The successful transfer of methyl group assignments to glycosylated Fc fragments of mAbs produced in CHO cells at natural isotopic abundance represents a major step forward for the structural characterization of therapeutic monoclonal antibodies under native-like conditions. This assignment enables the precise localisation of structural modifications, such as those induced by stress, degradation, or interaction with antigens or receptors, directly within the 3D structure of the Fc region, without the need for isotopic labelling. Such capability will be particularly valuable in the context of quality control and comparability assessments during mAb development. By overlaying methyl NMR spectra from different production batches, it becomes possible to detect subtle conformational changes and identify specific residues affected by post-translational modifications or process-related variations. This atomic-level resolution will provide a powerful analytical tool to ensure batch consistency, monitor structural integrity, and support regulatory compliance throughout the development of therapeutic antibodies.

### Studying the effects of methionine oxidation on the Fc fragment by NMR

The methyl group assignment established for the glycosylated Fc fragment produced in CHO cells provides a powerful tool for probing structural changes directly on therapeutic mAb batches. This approach is particularly relevant during the pharmaceutical development of new antibodies, where standard stress protocols are applied to simulate harsh storage conditions and assess molecular stability. Notably, methionine oxidation was observed following stress exposure and was found to impact pharmacokinetics (Stracke et al. 2014) and complement-dependent cytotoxicity activity (Mo et al. 2016; unpublished internal data). As reported in the literature, oxidation is among the most frequent chemical modifications affecting mAbs during production and storage and can alter biological activity (Nowak et al. 2017; Torosantucci et al. 2014). Moreover, NMR has already proven to be a powerful tool for monitoring the effect of methionine oxidation on mAb batches. For instance, 2D methyl fingerprints of model NISTmAb were analysed following complete methionine oxidation, revealing not only strong differences in the methionine signals region, but also chemical shift changes in other regions of the spectrum (Solomon et al. 2023). This study hypothesized that oxidation could affect the local structure surrounding the oxidized methionines; however, no site-specific information could be obtained due to the lack of methyl group assignment. On the Fc region, 2D ^1^H-^15^N spectra were compared upon methionine oxidation using non-glycosylated Fc fragments produced in *E. coli* and uniformly isotopically labelled with ^2^H, ^15^N, and ^13^C (Liu et al. 2008). The methyl group assignment of the Fc fragment from a mAb produced in CHO cells at natural abundance now provides a straightforward and efficient approach to identify amino acids affected by methionine oxidation, by overlaying 2D spectral fingerprints and detecting shifts in methyl signals, directly on mAb batches from biotherapeutic production pipelines.

To investigate the impact of methionine oxidation on the higher-order structure (HOS) of the mAb, particularly within the Fc region, partial chemical oxidation was induced by incubating overnight the anti-LAMP1 mAb with 2.4% (v/v) tBHP. Following mAb cleavage, SEC-MS intact mass spectrometry analysis of the fragments revealed near-complete oxidation of the two methionines of the Fc, M252 and M428, and no oxidation in the Fab fragment (Fig. S8). A 2D methyl NMR spectrum of the oxidized Fc was acquired and compared to the native Fc spectrum (Fig. 5).

**Fig. 5.**
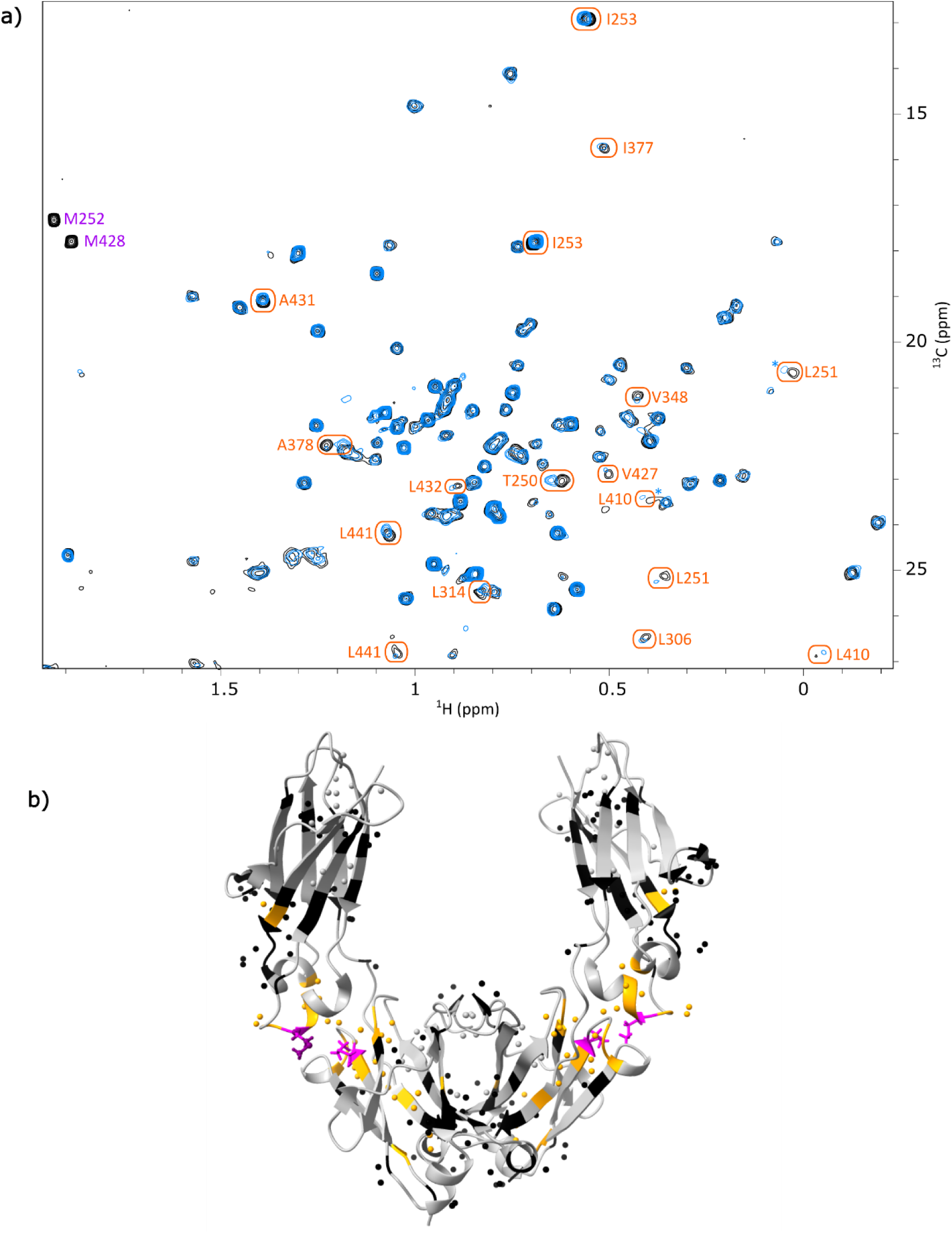
**a)** Superimposition of ^1^H-^13^C ALSOFAST methyl spectra of the glycosylated Fc fragment from anti-LAMP1 mAb before (in black) and after (in blue) methionine oxidation. Orange circles indicate assigned methyl resonances for which CSP values are above 0.01 ppm. Upon oxidation, methionine signals shift outside of the spectrum window. Asterisks indicate signals for which the contour level was multiplied by a factor of 2. **b)** Impact of methionine oxidation shown on the structure of a glycosylated Fc fragment (PBD: 3JII). Methyl groups are represented by spheres. Non-methylated amino acids and non-assigned methylated amino acids are in grey, assigned methylated amino acids are in black when CSP < 0.01 ppm and in orange when CSP ≥ 0.01 ppm, and methionines are in purple. For amino acids with two methyl groups (isoleucines, leucines, and valines), average CSPs were used when both signals were assigned

The signals corresponding to M252 and M428 were no longer visible, having shifted from around δ_H_ = 2.0 ppm / δ_C_ = 17.0 ppm to around δ_H_ = 2.6 ppm / δ_C_ = 39.9 ppm, outside the typical methyl spectral window. After transferring the assignments from the native to the oxidized Fc spectrum, chemical shift perturbations were calculated for all assigned methyl signals (Fig. S9). Although some assignments could not be transferred due to signal overlap or low intensity, CSPs were successfully calculated for 78% of methylated residues. While most CSPs were rather weak, several signals showed significant shifts. The corresponding amino acids are shown in orange on the Fc structure (Fig. 5). These residues were found to be located around the oxidized methionines, in both C_H_2 and C_H_3 domains, indicating localised structural changes near the oxidized M252 and M428. No long-range effects were observed, confirming that oxidation-induced changes are spatially restricted. This newly developed NMR approach enabled the straightforward identification of residues impacted by this stress, simply by overlaying 2D methyl spectra acquired directly on the Fc fragment of industrial mAb batches at natural abundance.

## Conclusions

By integrating cell-free expression, selective isotope labelling, and the acquisition of high-quality 2D and 3D NMR spectra, we successfully assigned 94% of methyl group resonances of a non-glycosylated Fc fragment, without requiring protein perdeuteration. This assignment was then transferred to a 2D methyl spectrum of a glycosylated Fc fragment obtained via enzymatic cleavage of a monoclonal antibody produced in CHO cells at natural isotopic abundance. The resulting spectral fingerprint, assigned at 83%, provides a solid foundation for atomic-resolution NMR studies of the Fc structure in IgG1 antibodies produced in eukaryotic systems. This work demonstrates that methyl NMR fingerprinting, combined with robust assignment strategies, enables rapid and precise identification of structural modifications in mAbs produced at natural abundance. The relevance of this approach was further highlighted by the identification of residues affected by methionine oxidation, a common degradation pathway observed under standard stress conditions. The method is broadly applicable to standard IgG1 biomolecules and can be routinely used to monitor batch-to-batch consistency, detect degradation, and support quality control throughout therapeutic antibody development and manufacturing. When combined with existing methyl group assignments of Fab fragments produced in vitro (Ghasriani et al. 2022; Gagné et al. 2024; Sarker and Aubin 2024; Henot et al. 2025), this strategy opens the door to comprehensive structural investigations of full-length IgG1 antibodies using standard 2D methyl NMR spectroscopy. Overall, this approach paves the way for atomic-resolution structural characterization and quality control of therapeutic mAbs during development phases. By enabling the detection of subtle conformational changes and stress-induced modifications directly on CHO-produced antibody batches, methyl NMR fingerprinting emerges as a powerful tool for comparability studies, stability assessments, and process optimization in biopharmaceutical development.

## Acknowledgements

The authors thank Rida Awad, Séverine Beauvisage, Claire Borel, Dawid Bugnazet, Ronan Crepin, Arthur Giraud, Lionel Imbert, Louis Joos, Alessandro Masiero, Alexey Rak, and Nicola Salvi for advice, support and stimulating discussions. This work is supported by the French National Research Agency in the framework of the “Investissements d’avenir” program (ANR-15-IDEX-02), NMR4mAbs project n° ANR-22-CE29-0024-01, the CEA/CNRS/UGA/SANOFI collaborative research program 2023-0666/C44988/231021, and the French Proteomic Infrastructure (ANR-10-INBS-08 & ANR-24-INBS-0015). This work used the high field NMR and Cell-Free facilities at the Grenoble Instruct-ERIC Center (ISBG; UAR 3518 CNRS-CEA-UGA-EMBL) within the Grenoble Partnership for Structural Biology (PSB). Platform access was supported by FRISBI (ANR-10-INBS-05-02) and GRAL, a project of the University Grenoble Alpes graduate school (Ecoles Universitaires de Recherche) CBH-EUR-GS (ANR-17-EURE-0003). IBS acknowledges integration into the Interdisciplinary Research Institute of Grenoble (IRIG, CEA). S.N. and B.V. acknowledge ANRT and Sanofi for PhD funding.

## Author contributions

B.V. and S.N. prepared the samples. B.V., S.N., F.H., C.D., O.F. and J.B. collected and analysed NMR data. S.N., O.H.-A., S.Ci., and S.Cl. collected and analysed mass spectrometry data. B.V., S.N., F.H., O.F. and J.B. wrote the manuscript. B.V. and S.N. prepared the figures. All authors reviewed the manuscript and approved the final version.

## Data availability

Chemical shifts and Bruker raw data files will be deposited in the BMRB data bank with entry numbers 53464 and 53465.

## Competing Interests

B.V., S.N., F.H., C.D., S.Cl., and O.F. are employees of Sanofi and may hold shares and/or stock options in the company.

## References

1. Arbogast LW, Delaglio F, Schiel JE, Marino JP (2017) Multivariate analysis of two-dimensional 1H, 13C methyl NMR spectra of monoclonal antibody therapeutics to facilitate assessment of higher order structure. Anal Chem 89(21):11839–11845. 10.1021/acs.analchem.7b03571

2. Arbogast LW, Delaglio F, Tolman JR, Marino JP (2018) Selective suppression of excipient signals in 2D ^1^H–^13^C methyl spectra of biopharmaceutical products. J Biomol NMR 72:149–161. 10.1007/s10858-018-0214-1

3. Brinson RG, Marino JP, Delaglio F, Arbogast LW et al. (2018) Enabling adoption of 2D-NMR for the higher order structure assessment of monoclonal antibody therapeutics. mAbs 11(1):94–105. 10.1080/19420862.2018.1544454

4. Cerofolini L, Ravera E, Fischer C, Trovato A, Sacco F, Palinsky W, Angiuoni G, Fragai M, Baroni F (2023) Integration of NMR spectroscopy in an analytical workflow to evaluate the effects of oxidative stress on abituzumab: beyond the fingerprint of mAbs. Anal Chem 95(24):9199–9206. 10.1021/acs.analchem.3c00317

5. Delaglio F, Grzesiek S, Vuister GW, Zhu G, Pfeifer J, Bax A (1995) NMRPipe: a multidimensional spectral processing system based on UNIX pipes. J Biomol NMR 6:277–293. 10.1007/BF00197809

6. Favier A, Brutscher B (2019) NMRlib: user-friendly pulse sequence tools for Bruker NMR spectrometers. J Biomol NMR 73:199–211. 10.1007/s10858-019-00249-1

7. Gagné D, Aramini JM, Aubin Y (2024) Backbone and methyl side-chain resonance assignments of the single chain Fab fragment of trastuzumab. Biomol NMR assign, 18(2):119–128. 10.1007/s12104-024-10177-3

8. Gagné D, Sarker M, Gingras G, Hodgson DJ, Frahm G, Creskey M, Lorbetskie B, Bigelow S, Wang J, Zhang X, Johnston MJW, Lu H, Aubin Y (2023) Strategies for the production of isotopically labelled Fab fragments of therapeutic antibodies in Komagataella phaffii (Pichia pastoris) and Escherichia coli for NMR studies. PLoS One 18(11):e0294406. 10.1371/journal.pone.0294406

9. Gans P, Boisbouvier J, Ayala I, Hamelin O (2013) Process for the specific isotopic labeling of methyl groups of Val, Leu and Ile. Patent number: 9708228.

10. Ghasriani H, Ahmadi S, Hodgson DJ, Aubin Y (2022) Backbone and side-chain resonance assignments of the NISTmAb-scFv and antigen-binding study. Biomol NMR assign 16(2):391–398. 10.1007/s12104-022-10109-z

11. Giraud A, Imbert L, Favier A, Henot F, Duffieux F, Samson C, Frances O, Crublet E, Boisbouvier J (2024) Enabling site–specific NMR investigations of therapeutic Fab using a cell–free based isotopic labeling approach: application to anti–LAMP1 Fab. J Biomol NMR. 78:73–86. 10.1007/s10858-023-00433-4

12. Goto NK, Gardner KH, Mueller GA, Willis RC, Kay LE (1999) A robust and cost-effective method for the production of Val, Leu, Ile (delta 1) methyl-protonated 15N-, 13C-, 2H-labeled proteins. J Biomol NMR 13:369–374. 10.1023/a:1008393201236

13. Henot H, Vibert B, Giraud A, Dbira S, Imbert L, Favier A, Güntert P, Clavier S, Crublet E, Doyen C, Boisbouvier J, Frances O (accepted) A fast and efficient strategy for the NMR assignment of Fab methyl groups. J. Biomol. NMR. 10.1101/2025.10.21.682867

14. Imbert L, Lenoir-Capello R, Crublet E, Vallet A, Awad R, Ayala I, Juillan-Binard C, Mayerhofer H (2021) In vitro production of perdeuterated proteins in H_2_O for biomolecular NMR studies. Structural Genomics 2199:127–149. 10.1007/978-1-0716-0892-0_8

15. Kay LE, Xu GY, Singer AU, Muhandiram DR, Formankay JD (1993) A Gradient-Enhanced HCCH-TOCSY Experiment for Recording Side-Chain ^1^H and ^13^C Correlations in H _2_O Samples of Proteins. J Magn Reson 101(3):333–337. 10.1006/jmrb.1993.1053

16. Kerfah R, Plevin MJ, Sounier R, Gans P, Boisbouvier J (2015) Methyl-specific isotopic labeling: a molecular tool box for solution NMR studies of large proteins. Curr Opin Struct Biol 32:113–122. 10.1016/j.sbi.2015.03.009

17. Kothari M, Wanjari A, Acharya S, Karwa V, Chavhan R, Kumar S, Kadu A, Patil R (2024) A Comprehensive Review of Monoclonal Antibodies in Modern Medicine: Tracing the Evolution of a Revolutionary. Therapeutic Approach Cureus 16(6):e61983. 10.7759/cureus.61983

18. Lescop E, Schanda P, Brutscher B (2007) A set of BEST triple-resonance experiments for time-optimized protein resonance assignment. J Magn Res 187(1):163–169. 10.1016/j.jmr.2007.04.002

19. Liu D, Cocco M, Rosenfied R, Lewis J, Ren D, Li L, Remmele RLJ, Brems DN (2007a) Assignment of backbone ^1^H, ^13^C and 15N resonances of human IgG1 Fc (51.4 kDa). Biomol NMR Assign 1:233–235. 10.1007/s12104-007-9065-5

20. Liu D, Cocco M, Matsumura M, Ren D, Becker B, Remmele RLJ, Brems DN (2007b) Assignment of ^1^H, ^13^C and ^15^N resonances of the reduced human IgG1 C_H_3 domain. Biomol NMR Assign 1:93–94 10.1007/s12104-007-9026-z

21. Liu D, Ren D, Huang H, Dankberg J, Rosenfeld R, Cocco MJ, Li L, Brems DN, Remmele RLJ (2008) Structure and Stability Changes of Human IgG1 Fc as a Consequence of Methionine Oxidation. Biochemistry 47(18):5088–5100. https://pubs.acs.org/doi/10.1021/bi702238b

22. Marino JP, Brinson RG, Hudgens JW, Ladner JE, Gallagher DT, Gallagher ES, Arbogast LW, Huang RYC (2015) Emerging technologies to assess the higher order structure of monoclonal antibodies. ACS Symposium Series 1202(2):17–43. 10.1021/bk-2015-1202.ch002

23. Martins AC, Oshiro MY, Schiavon BN, de Jesus GA, de la Torre BG, Albericio F (2025) Monoclonal Antibodies (mAbs) and Proteins: The Biologic Drugs Approved by the Food and Drug Administration (FDA) in 2024. Biomedicines 13(8):1962. 10.3390/biomedicines13081962

24. Mo J, Yan Q, So CK, Soden T, Lewis MJ, Hu P (2016) Understanding the Impact of Methionine Oxidation on the Biological Functions of IgG1 Antibodies Using Hydrogen/Deuterium Exchange Mass Spectrometry. Anal Chem 88(19):9495–9502. 10.1021/acs.analchem.6b01958

25. Mueller L (2008) Alternate HMQC experiments for recording HN and HC-correlation spectra in proteins at high throughput. J Biomol NMR 42:129–137. 10.1007/s10858-008-9270-2

26. Nowak C, Cheung JK, Dellatore SM, Katiyar A, Bhat R, Sun J, Ponniah G, Neill A, Mason B, Beck A, Liu H (2017) Forced degradation of recombinant monoclonal antibodies: a practical guide. mAbs 9(8):74–85. 10.1080/19420862.2017.1368602

27. Pervushin K, Riek R, Wider G, Wüthrich K (1997) Attenuated T2 relaxation by mutual cancellation of dipole-dipole coupling and chemical shift anisotropy indicates an avenue to NMR structures of very large biological macromolecules in solution. Proc Natl Acad Sci U S A. 94:12366–12371. 10.1073/pnas.94.23.12366

28. Rößler P, Mathieu D, Gossert AD (2020) Enabling NMR Studies of High Molecular Weight Systems Without the Need for Deuteration: The XL-ALSOFAST Experiment with Delayed Decoupling. Angew Chem Int Ed Engl 59(43):19329–19337. 10.1002/anie.202007715

29. Sarker M, Aubin Y (2024) Backbone and methyl side-chain resonance assignments of the Fab fragment of adalimumab. Biomol NMR assign 18(2):187–192. 10.1007/s12104-024-10187-1

30. Schanda P, Brutscher B (2005) Very Fast Two-Dimensional NMR Spectroscopy for Real-Time Investigation of Dynamic Events in Proteins on the Time Scale of Seconds. J Am Chem Soc 127(22):8014–8015. 10.1021/ja051306e

31. Solomon TL, Delaglio F, Giddens JP, Marino JP, Yu YB, Taraban MB, Brinson RG (2023) Correlated analytical and functional evaluation of higher order structure perturbations from oxidation of NISTmAb. mAbs 15(1) 10.1080/19420862.2022.2160227

32. Stracke J, Emrich T, Rueger P, Schlothauer T, Kling L, Knaupp A, Hertenberger H, Wolfert A, Spick C, Lau W, Drabner G, Reiff U, Koll H, Papadimitriou A (2014) A novel approach to investigate the effect of methionine oxidation on pharmacokinetic properties of therapeutic antibodies. Mabs 6(5):1229–42. 10.4161/mabs.29601

33. Torosantucci R, Schöneich S, Jiskoot W (2014) Oxidation of Therapeutic Proteins and Peptides: Structural and Biological Consequences. Pharm Res 3(31):541–553 10.1007/s11095-013-1199-9

34. de la Torre BG, Albericio F (2025) The Pharmaceutical Industry in 2024: An Analysis of the FDA Drug Approvals from the Perspective of Molecules. Molecules 30(3):482. 10.3390/molecules30030482

35. Tugarinov V, Kay LE (2004) An Isotope Labeling Strategy for Methyl TROSY Spectroscopy. J Biomol NMR 28:165–172. 10.1023/B:JNMR.0000013824.93994.1f

36. Tugarinov V, Kay LE (2005) Methyl groups as probes of structure and dynamics in NMR studies of high-molecular-weight proteins. Chembiochem 6(9):1567–1577. 10.1002/cbic.200500110

37. Tugarinov V, Kay LE, Ibraghimov I, Orekhov VY (2005) High resolution four-dimensional ^1^H-^13^C NOE spectroscopy using methyl-TROSY, sparse data acquisition, and multidimensional decomposition. J Am Chem Soc 127:2767–2775. 10.1021/ja044032o

38. Vranken WF, Boucher W, Stevens TJ, Fogh RH, Pajon A, Llinas M, Ulrich EL, Markley JL, Ionides J, Laue ED (2005) The CCPN data model for NMR spectroscopy: development of a software pipeline. Proteins 59:687–696. 10.1002/prot.20449

39. Yin G, Swartz JR (2004) Enhancing multiple disulfide bonded protein folding in a cell-free system. Biotechnol Bioeng 86(2):188–195. 10.1002/bit.10827

40. Yagi H, Zhang Y, Yagi-Utsumi M, Yamaguchi T, Iida S, Yamaguchi Y, Kato K (2015) Backbone 1H, 13 C, and 15 N resonance assignments of the fc fragment of human immunoglobulin G glycoprotein. Biomol NMR Assign 9(2):257–260. 10.1007/s12104-014-9586-7

