## Supplementary information for "Toward Site-Specific Characterization of Structural Perturbations on Glycosylated Fc Using NMR at Natural Abundance"

[*] Corresponding authors

### Supplementary information

**Sequence** of the non-glycosylated Fc fragment produced in a cell-free system (M220 - H464)

**Sequence** of the glycosylated Fc fragment obtained from a mAb produced in CHO cells (H224 - H446)

**Table S1** List of labelled samples and NMR experiments acquired on the non-glycosylated Fc

**Fig. S1** Quality control of an Fc sample produced in a cell-free system at natural abundance

**Fig. S2** Assigned NH backbone resonances of the Fc fragment produced in a cell-free system

**Fig. S3** Valines assignment strategy for the non-glycosylated Fc fragment

**Fig. S4** Schematic representation of the methyl group resonances assignment of the Fc fragment

**Fig. S5** Overlay of the methyl spectra of the non-glycosylated Fc fragment produced in a cell-free system and the glycosylated Fc fragment from anti-LAMP1 mAb produced in CHO cells

**Fig. S6** Overlay of the methyl spectra of the glycosylated and the deglycosylated Fc fragments

**Fig. S7** CSPs measured between the assignment of non-glycosylated and glycosylated Fc fragments

**Fig. S8** SEC-native MS analysis of anti-LAMP1's $F(ab')_2$ Fc subunits before and after oxidation

**Fig. S9** CSPs measured upon methionine oxidation of the glycosylated Fc fragment

**Sequence of the non-glycosylated Fc fragment produced in a cell-free system (M220 - H464)**

MDKTHTCPPCPAPELLGGPSVFLFPPKPKDTLMISRTPEVTCVVVDVSHEDPEVKFNWYVDGVEVHNAKTKPREEQ
YNSTYRVVSVLTVLHQDWLNGKEYKCKVSNKALPAPIEKTISKAKGQPREPQVYTLPPSRDELTKNQVSLTCLVKGFYP
SDIAVEWESNGQPENNYKTTPPVLDSDGSFFLYSKLTVDKSRWQQGNVFSCSVMHEALHNHYTQKSLSLSPGENLY
FQGPGGGSHHHHHH

**Sequence of the glycosylated Fc fragment obtained from a mAb produced in CHO cells (H224 - H446)**

HTCPPCPAPELLGGPSVFLFPPKPKDTLMISRTPEVTCVVVDVSHEDPEVKFNWYVDGVEVHNAKTKPREEQYNSTY
RVVSVLTVLHQDWLNGKEYKCKVSNKALPAPIEKTISKAKGQPREPQVYTLPPSRDELTKNQVSLTCLVKGFYPSDIAV
EWESNGQPENNYKTTPPVLDSDGSFFLYSKLTVDKSRWQQGNVFSCSVMHEALHNHYTQKSLSLSPG

**Table S1**: List of labelled samples and NMR experiments acquired on the non-glycosylated Fc fragment to perform methyl group resonances assignment

| Sample labelling | C (µM) | Experiment | Time (h) | Magn. Field (MHz) | NS | d1 (s) | SW F1/F2 (ppm) | TD F1/F2 |
|---|---|---|---|---|---|---|---|---|
| U-[$^{15}N$, $^{13}C$] | 300 | $^{1}H$-$^{15}N$ BEST-TROSY-HSQC | 0.5 | 850 | 12 | 0.2 | 36 | 434 |
| | | BEST-TROSY-HNCO | 15.5 | 600 | 8 | 0.2 | 15/36 | 128/148 |
| | | BEST-TROSY-HNCA | 62.3 | 850 | 32 | 0.2 | 30/36 | 128/148 |
| U-[$^{15}N$, $^{13}C$] | 700 | $^{1}H$-$^{13}C$ CT-SOFAST-HMQC | 0.2 | 600 | 32 | 0.2 | 15 | 104 |
| | | hCCH TOCSY | 89.6 | 600 | 8 | 1.0 | 75/75 | 188/188 |
| A, I-$\delta_1$, M, T, V$\gamma_1$/$\gamma_2$: $^{13}CH_3$ | 1600 | CCH HMQC-NOESY-HMQC | 62.3 | 950 | 12 | 0.9 | 15/15 | 128/128 |
| L-$\delta_1$, L-$\delta_2$: $^{13}CH_3$ | 785 | CCH HMQC-NOESY-HMQC | 96.1 | 700 | 136 | 0.8 | 10/10 | 50/50 |
| | | $^{1}H$-$^{13}C$ CT-SOFAST-HMQC | 1.6 | 600 | 128 | 0.5 | 10 | 72 |
| | | $^{1}H$-$^{13}C$ SOFAST-HMQC | 2.2 | 600 | 64 | 0.5 | 10 | 212 |
| V-$\gamma_1$,$\gamma_2$: $^{13}CH_3$ | 225 | CCH HMQC-NOESY-HMQC | 113.2 | 950 | 80 | 0.8 | 10/10 | 70/70 |
| M, V-$\gamma_1$/$\gamma_2$: $^{13}CH_3$ | 160 | $^{1}H$-$^{13}C$ SOFAST-HMQC | 1.2 | 700 | 48 | 0.5 | 9 | 158 |
| I-$\delta_1$, T: $^{13}CH_3$ | 164 | $^{1}H$-$^{13}C$ SOFAST-HMQC | 0.9 | 700 | 24 | 0.5 | 14 | 246 |
| A: $^{13}CH_3$ | 232 | $^{1}H$-$^{13}C$ SOFAST-HMQC | 0.9 | 700 | 24 | 0.5 | 5 | 88 |

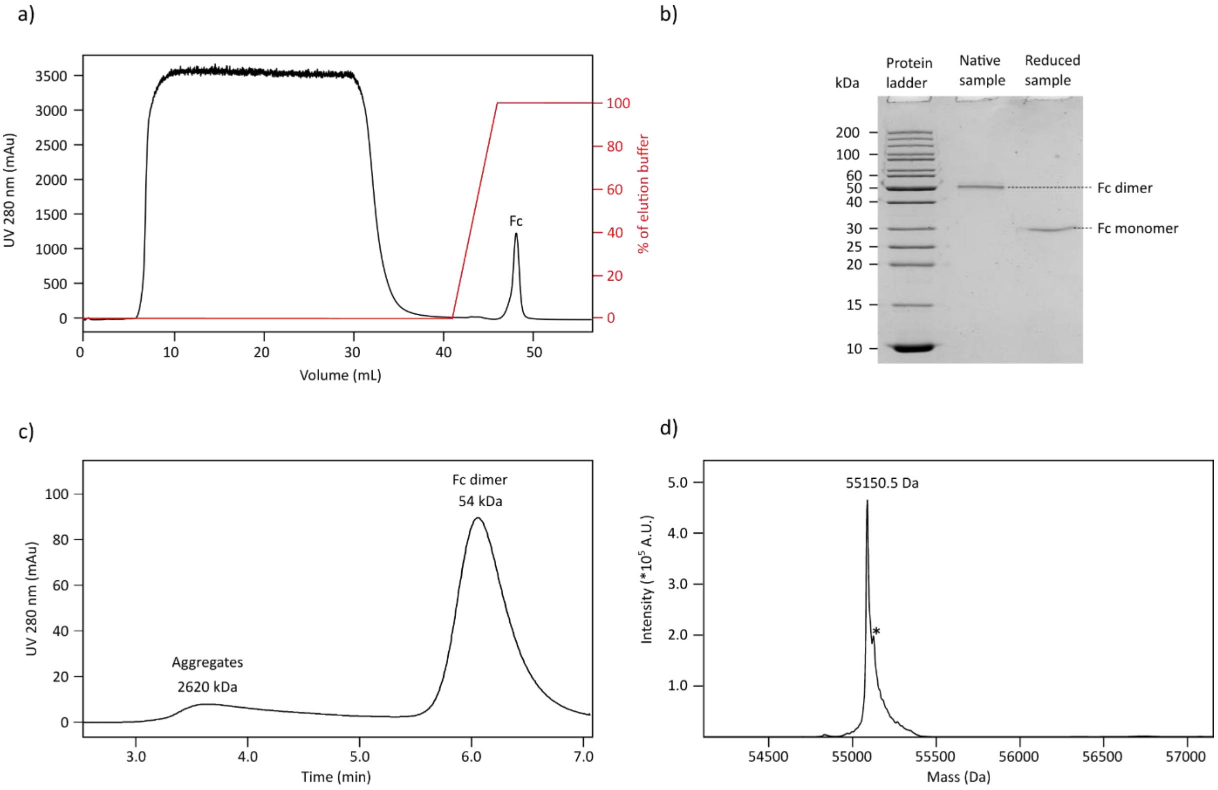


**Fig. S1** Quality control of an Fc sample produced in a cell-free system at natural abundance. **a)** Purification of 6 mL of a cell-free reaction mixture on a 1 mL CaptureSelect FcXP affinity chromatography column. **b)** 15% acrylamide SDS-PAGE gel. 0.2 µg of purified Fc sample were loaded in the second (native condition) and third (reductive condition) wells. **c)** SEC-MALS results obtained for the injection of 10 µg of an Fc sample on a Superdex 200 Increase 5/150 (Cytiva) column. **d)** Deconvoluted mass spectrum of 3 µg of a purified Fc sample following analysis in RPLC-MS on the BioAccord LC-MS mass spectrometer (Waters). The peak annotated with an asterisk corresponds to the acetonitrile adduct

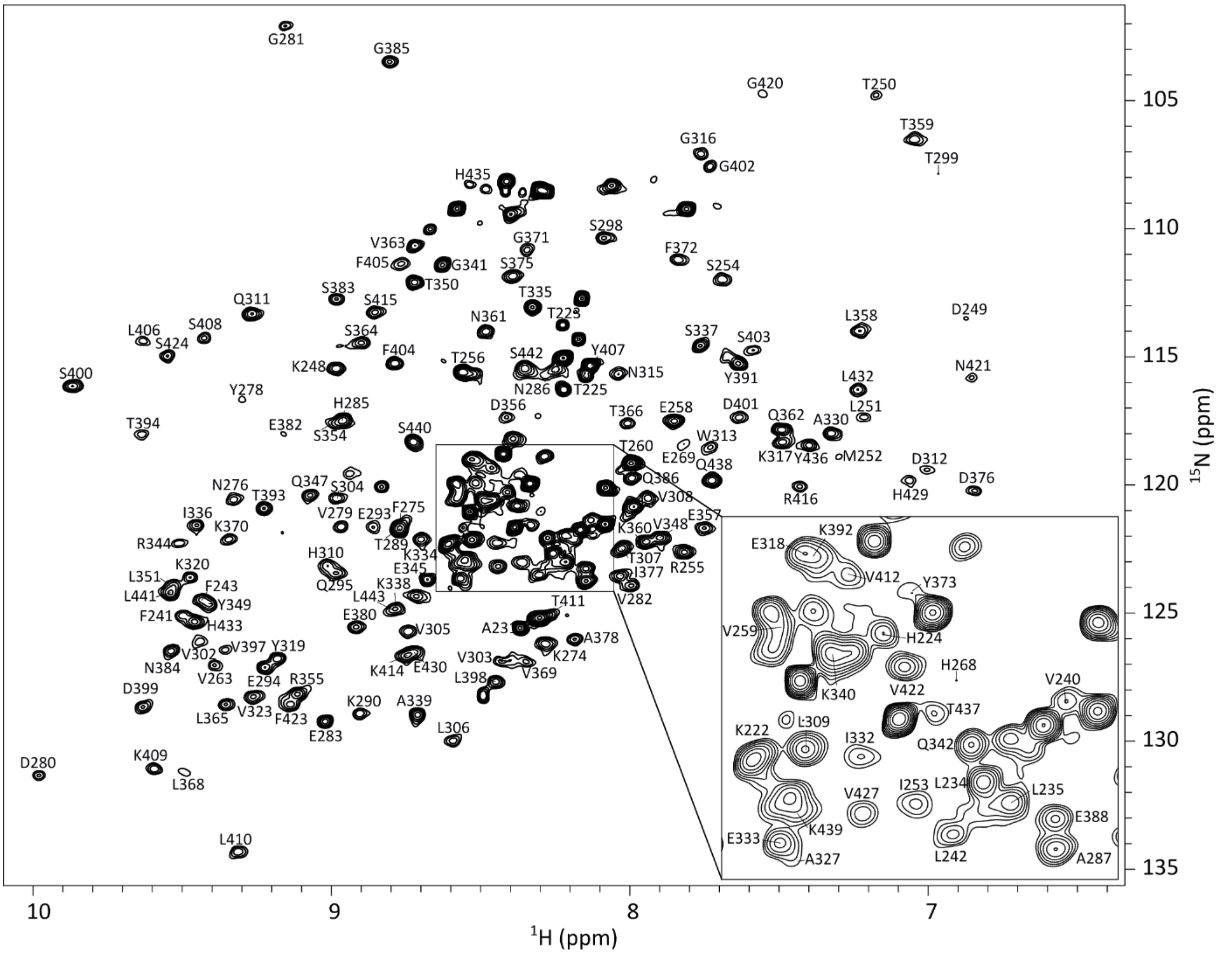


**Fig. S2** Assigned 2D $^1$H-$^{15}$N BEST-TROSY-HSQC spectrum acquired at 323 K on a U-[$^{15}$N, $^{13}$C] Fc fragment produced using a cell-free system. Assigned resonances are annotated with the corresponding amino acid

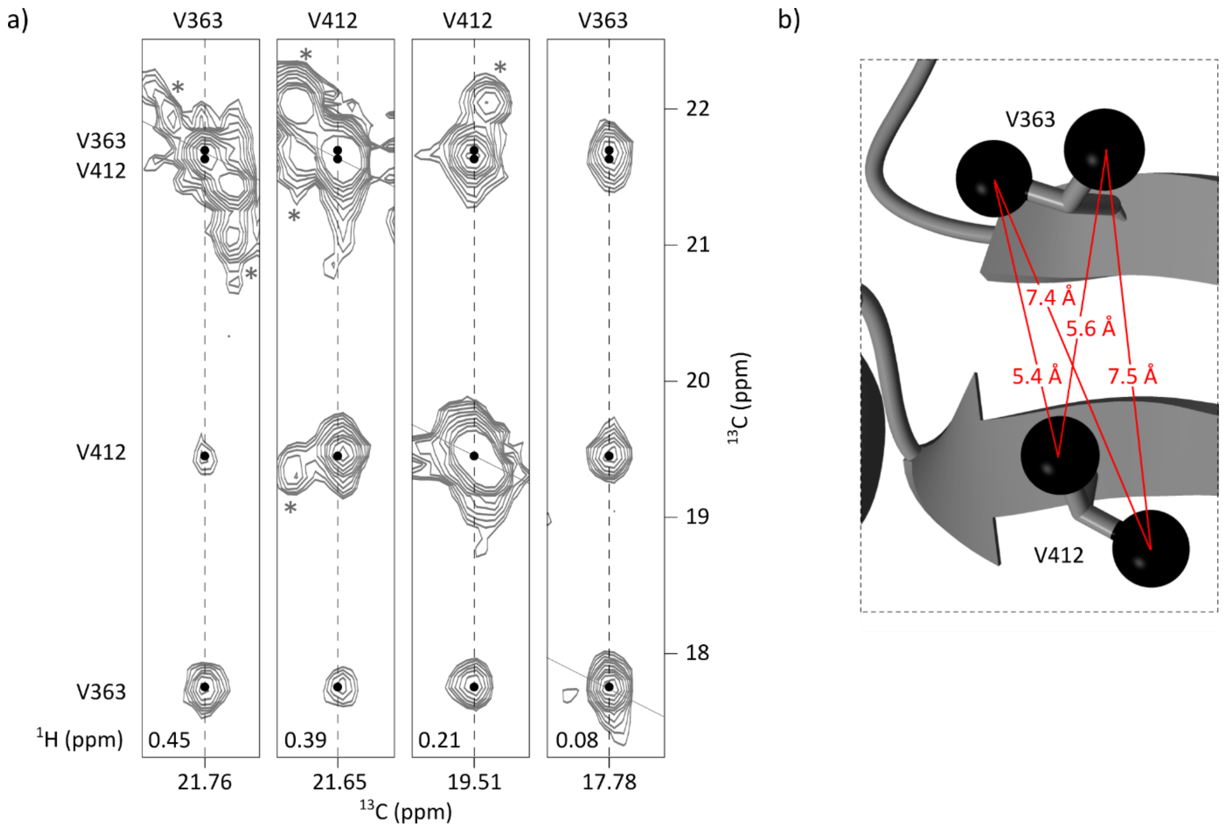


**Fig. S3** Valines assignment strategy for the non-glycosylated Fc fragment. **a)** 2D extracts from a 3D HMQC-NOESY-HMQC experiment acquired on an Fc fragment labelled on Val-[U-$^{2}$H; [$^{13}$C$^{1}$H$_3$]$^{\gamma1}$; [$^{13}$C$^{1}$H$_3$]$^{\gamma2}$] showing NOE cross-peaks between V363-$\gamma_1$, V363-$\gamma_2$, V412-$\gamma_1$, and V412-$\gamma_2$. Asterisks indicate signals outside of the considered plans. **b)** Zoom on V412 and V363 extracted from the 3D structure of non-glycosylated Fc fragment (PBD 3JII) showing Val-$\gamma_1$ and Val-$\gamma_2$ methyl groups

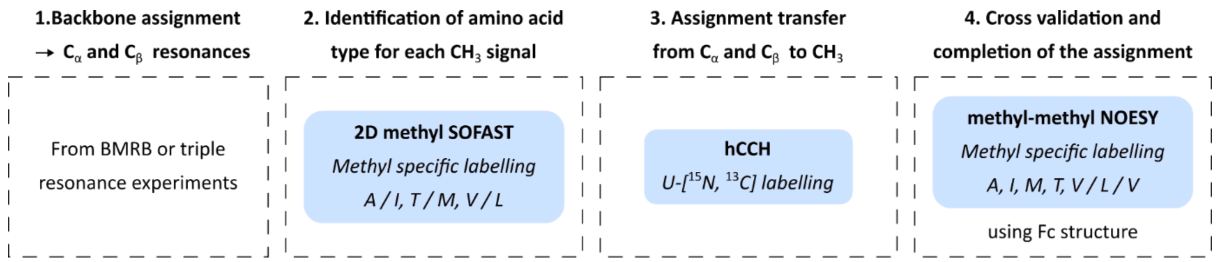


**Fig. S4** Schematic representation of the strategy used to assign the methyl group resonances of the non-glycosylated Fc fragment produced using a cell-free system

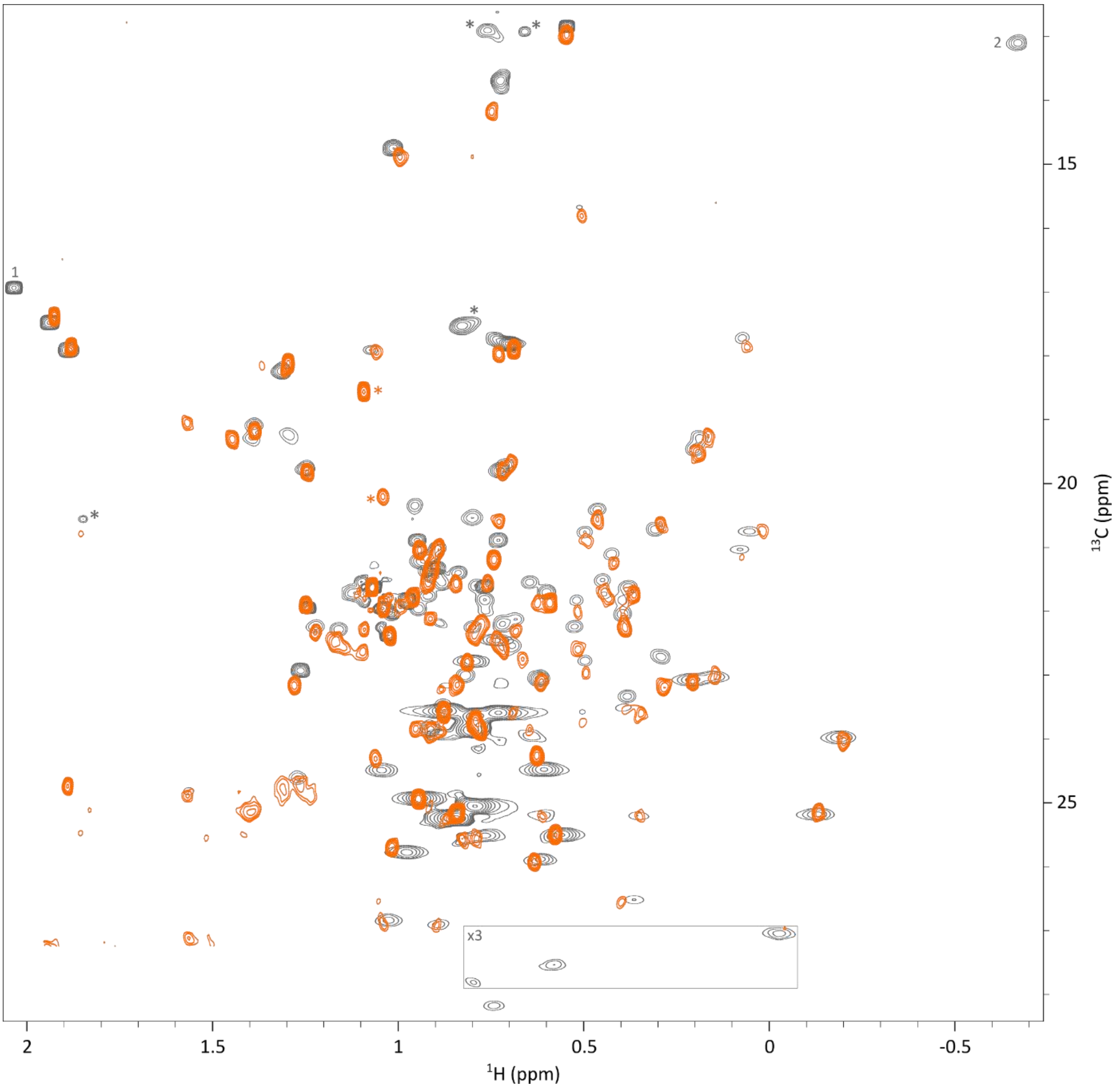


**Fig. S5** Overlay of $^{1}$H-$^{13}$C methyl spectra of the non-glycosylated Fc fragment produced in a cell-free system (grey, corresponding to Fig. 2) and the glycosylated Fc fragment from anti-LAMP1 mAb produced in CHO cells (orange, corresponding to Fig. 4 a). Signals marked with a grey asterisk correspond to impurities, and the two signals annotated with an orange asterisk correspond to methyl groups of N-acetylglucosamine (GlcNAc) moieties from the glycans. The contour level of grey signals within the grey rectangle was increased threefold. The grey signal labelled “1” corresponds to M220, which is absent from the sequence of the glycosylated Fc fragment from anti-LAMP1 produced in CHO cells and therefore not visible in the corresponding spectrum. The grey signal labelled “2” corresponds to I377-$\delta_1$, this signal is not observed in the spectrum of the glycosylated Fc fragment from anti-LAMP1 produced in CHO cells, due to the proton excitation bandwidth of the SOFAST-methyl-TROSY experiment

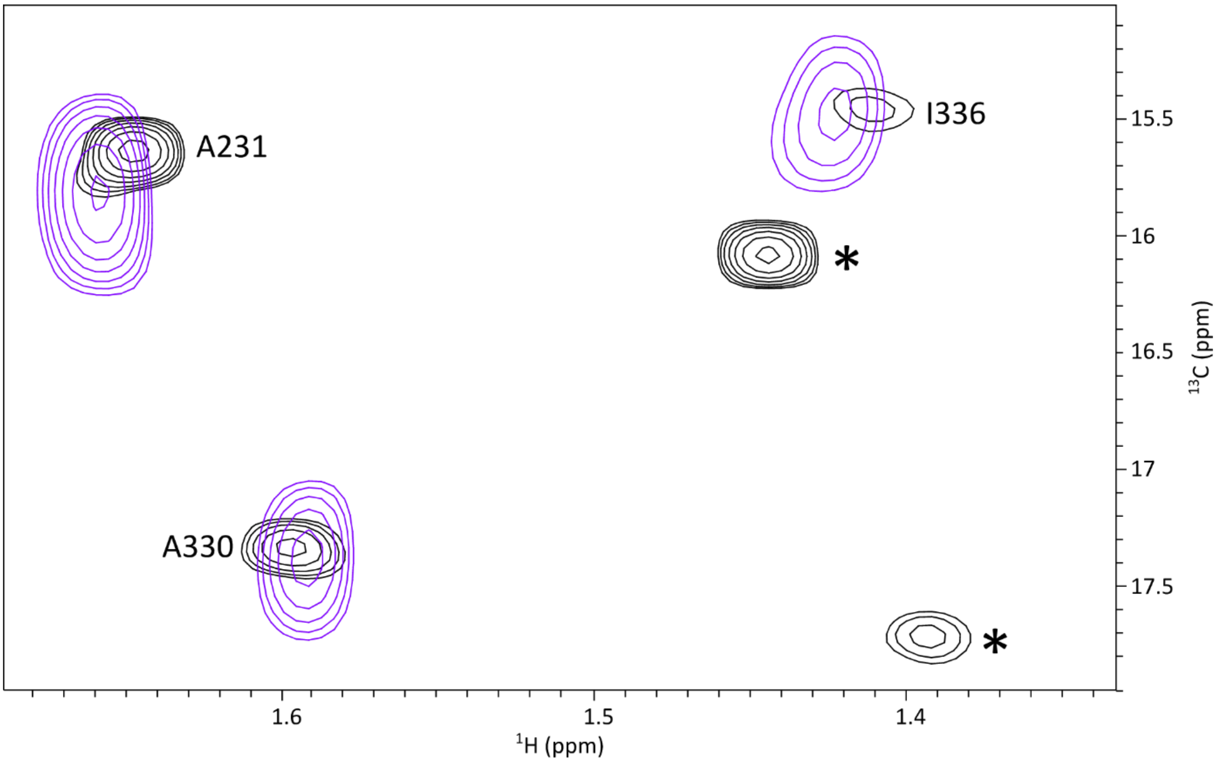


**Fig. S6** Overlay of 2D $^{1}H$-$^{13}C$ methyl spectra of the glycosylated Fc fragment (black) and the deglycosylated Fc fragment (purple). Methyl signals from GlcNAc residues of the glycans are marked with an asterisk. Deglycosylation of the Fc fragment was carried out by incubating a 1.2 g/L solution of glycosylated Fc with PNGase F (Genovis) for 60 min at 37 °C

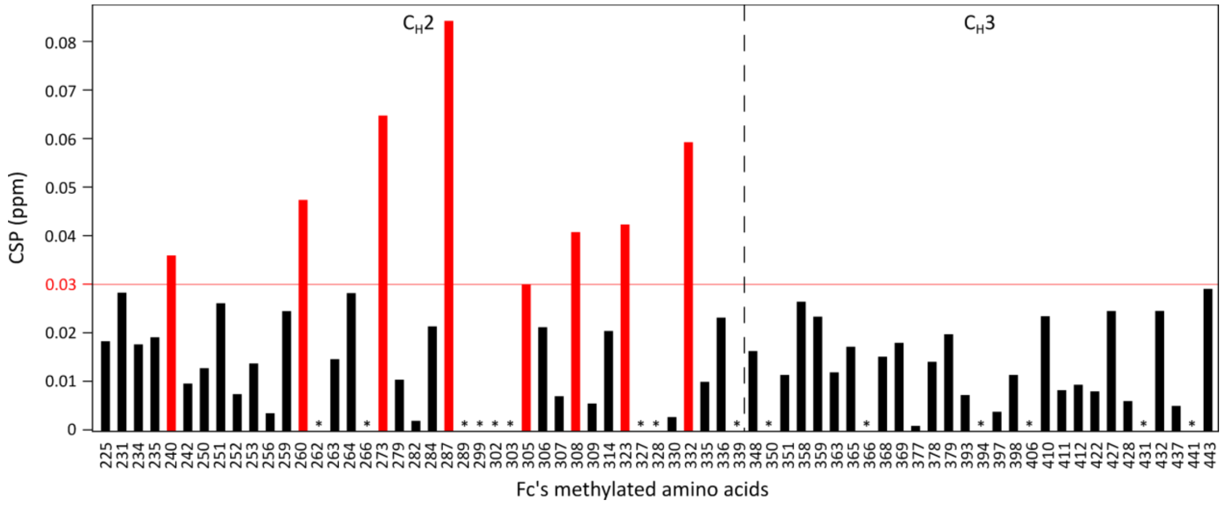


**Fig. S7** Chemical shift differences measured between the assignment of the non-glycosylated Fc fragment produced in cell-free and the glycosylated Fc fragment from anti-LAMP1 mAb produced in CHO cells. CSPs were calculated using the following formula: $CSP = \sqrt{\Delta\delta_H^2 + \left(\frac{\Delta\delta_C}{4}\right)^2}$. For isoleucines, leucines, and valines, the indicated CSP value is the mean of CSP values of the two methyl groups when both were assigned. CSP values above the threshold of 0.03 ppm are shown in red. Asterisks indicate methylated amino acids with no assigned resonance in at least one of the two spectra

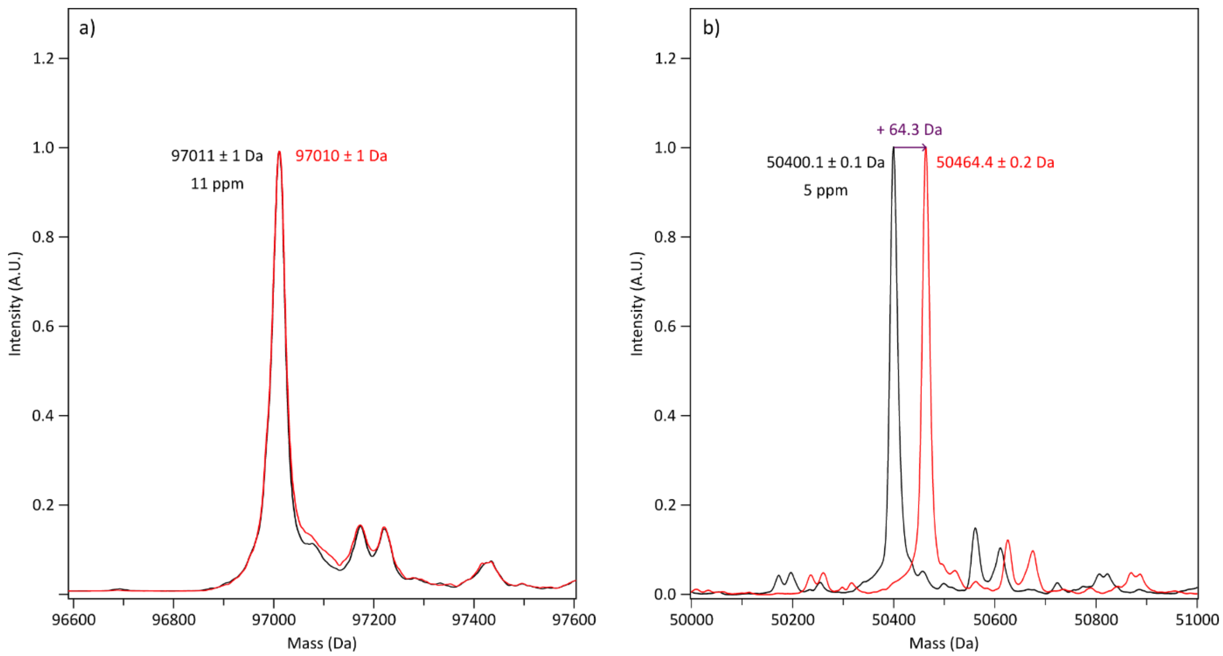


**Fig. S8**: SEC-MS analysis in non-denaturing conditions of anti-LAMP1 following FabRICATOR (IdeS) limited digestion (CPAPELLG/GPSV), resulting in (**a**) F(ab')$_2$ and (**b**) Fc subunits. Mass spectra of the non-oxidized (black trace) and oxidized (red trace) samples were overlayed. Following oxidative stress, the Fc displayed a mass increase of 64 Da, corresponding to the oxidation of four methionines: M252 and M428 of both chains

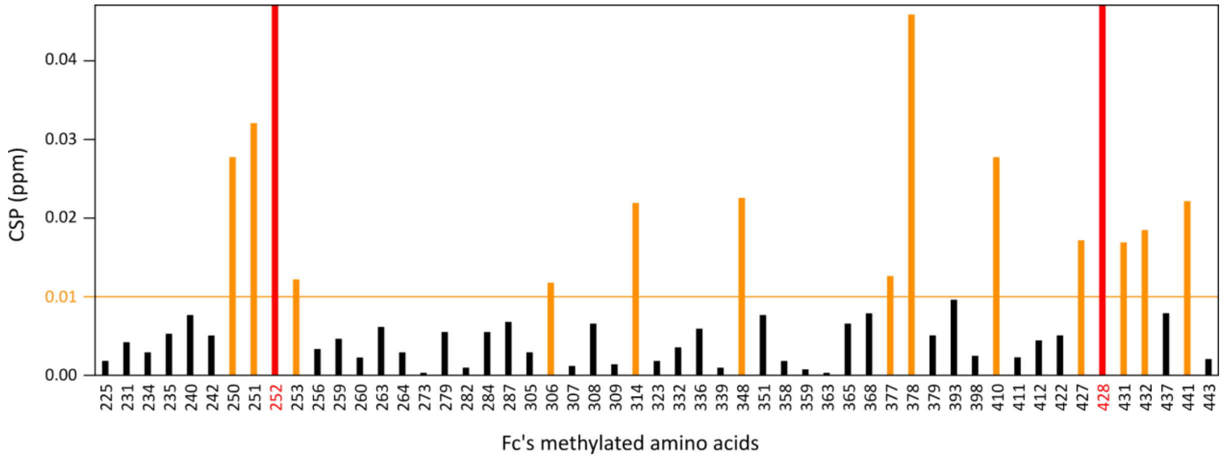


**Fig. S9** Chemical shift perturbations measured upon methionine oxidation of the glycosylated Fc fragment from anti-LAMP1 produced in CHO cells. CSP were calculated using the following formula: $CSP = \sqrt{\Delta\delta_H^2 + \left(\frac{\Delta\delta_C}{4}\right)^2}$ and the mean of CSP values was calculated for isoleucines, leucines, and valines when both methyl groups were assigned. CSP values above the threshold of 0.01 ppm are shown in orange and methionines are indicated with red bars
